# Optimized Data-Independent Acquisition Approach for Proteomic Analysis at Single-Cell Level

**DOI:** 10.1101/2021.08.19.457000

**Authors:** Yuefan Wang, Tung-Shing Mamie Lih, Lijun Chen, Yuanwei Xu, Morgan D. Kuczler, Liwei Cao, Kenneth J. Pienta, Sarah R. Amend, Hui Zhang

**Affiliations:** Department of Pathology, Johns Hopkins University,Baltimore, MD 21287, USA; The Brady Urological Institute, Johns Hopkins School of Medicine, 600 N. Wolfe Street, Baltimore, MD 21287, USA

## Abstract

Single-cell proteomic analysis provides valuable insights into cellular heterogeneity allowing the characterization of the cellular microenvironment which is difficult to accomplish in bulk proteomic analysis. Currently, single-cell proteomic studies utilize data-dependent acquisition (DDA) mass spectrometry (MS) coupled with a TMT labelled carrier channel. Due to the extremely imbalanced MS signals among the carrier channel and other TMT reporter ions, the quantification is compromised. Thus, data-independent acquisition (DIA)-MS should be considered as an alternative approach towards single-cell proteomic study since it generates reproducible quantitative data. However, there are limited reports on the optimal workflow for DIA-MS-based single-cell analysis. Herein, we report an optimized DIA workflow for single-cell proteomics using Orbitrap Lumos Tribrid instrument. We utilized a breast cancer cell line (MDA-MB-231) and induced drug resistant polyaneuploid cancer cells (PACCs) to evaluate our established workflow. We found that a short LC gradient was preferable for peptides extracted from single cell level with less than 2 ng sample amount. The total number of co-searching peptide precursors was also critical for protein and peptide identifications at nano- and sub-nano-gram levels. Post-translationally modified peptides could be identified from a nano-gram level of peptides. Using the optimized workflow, up to 1,500 protein groups were identified from a single PACC corresponding to 0.2 ng of peptides. Furthermore, about 200 peptides with phosphorylation, acetylation, and ubiquitination were identified from global DIA analysis of 100 cisplatin resistant PACCs (20 ng). Finally, we used this optimized DIA approach to compare the whole proteome of MDA-MB-231 parental cells and induced PACCs at a single-cell level. We found the single-cell level comparison could reflect real protein expression changes and identify the protein copy number. Our results demonstrate that the optimized DIA pipeline can serve as a reliable quantitative tool for single-cell as well as sub-nano-gram proteomic analysis.

## INTRODUCTION

Cells from the same living organism have a similar genomic background, which are eventually differentiated into diverse cell types in different tissues or organs via the expression of different genes to proteins leading to cellular heterogeneity. Although the rapid development of genomic and transcriptomic methods made it possible to analyze genomic and transcriptomic alterations of cellular heterogeneity at single-cell level^1,2^, the absence of protein amplification techniques hampered single-cell proteomic analysis. Originally, single-cell proteomic studies were limited to the detection of less than 15 targeted proteins from single mammalian cell by means of flow cytometry^3,4^, mass cytometry^5^, and single-cell western blotting^6^. After decades of development, mass spectrometry (MS), a primary tool for analyzing proteome and protein post-translational modifications (PTMs) from bulk samples, was applied to the first hypothesis-free mammalian single-cell proteomic analysis known as Single Cell ProtEomics by Mass Spectrometry (SCoPE-MS)^7^ in 2018. ScoPE-MS utilizes isobaric tandem mass tags (TMT) to label peptides from single cells along with a carrier sample containing highly excessive number of peptides to increase the detection of peptide fragment ions, especially for low abundant peptides, during tandem mass spectrometry analysis (MS/MS). Several works have followed the aforementioned concept and greatly expended the single-cell proteomics field^8^. However, TMT carrier-based methods have the limitations that the data quality and quantitation are highly dependent on the extremely imbalanced carrier ratio and instrument dynamic^9^.

Data independent acquisition (DIA)-MS is considered as a consistent proteomic analytical method that allows the fragmentation of all the precursor ions within selected isolation m/z range generating comprehensive MS/MS spectra^10^. DIA-MS can provide reproducible global quantitative data with minimal cost^10,11,12,13^. Various software tools have been developed^14,15^ to analyze DIA data that can be classified into spectral library-based approach^16^ and library-free approach ^17^. The spectral library-based DIA analysis is a peptide-centric method, which usually requires building spectral libraries either using corresponding DDA and/or DIA data from the same samples or using pre-built publicly available spectral libraries. However, the sample types and experimental conditions should be taken into consideration while building spectral libraries, especially when using external sources. Moreover, the spectral library size has direct impacts on DIA data search results^18^, thus, an inappropriate library size would compromise the identification results^19,20^. On the other hand, the library-free DIA analysis is a spectrum-centric method. There are several tools to conduct library-free analysis, including DIA-Umpire^17^ and directDIA embedded in Spectronaut^21^. Library-free approach detects or deconvolutes chromatographic features of precursor-fragment ion groups to generate pseudo-MS/MS spectra, which allows multiple DIA raw files to be processed together. While generating pseudo-MS/MS spectra from one or more DIA raw files, an internal spectral library is constructed like the pre-built library from DDA data/external sources. Therefore, the internal spectral library size could influent the DIA study. Nonetheless, library-free approach only relies on DIA data itself, thus, it is highly sample-specific compared to spectral library-based approach, which is more suitable for single cell global proteome identification.

In this study, we evaluated the performances of DIA-MS approach for the analysis at the nano and sub-nano-gram peptide levels using MDA-MB-231 cancer cells and drug resistant PACCs induced by platinum or docetaxel treatment^22^ to optimize the DIA-MS workflow for single-cell level proteomics. PACCs are a large cancer cell state with high genome content that are induced by stress and treatment. We evaluated the DIA performances in different liquid chromatography (LC), MS/MS, and data analysis settings. We found that a 15-minute short LC gradient and library-free approach via directDIA for data analysis allowed the identification of 3,260 and 1,530 proteins from 2 ng (corresponding to 10 PACCs) and 0.2 ng (corresponding to a single PACC) peptides with good reproducibility, respectively. Therefore, the results demonstrate that our optimized DIA pipeline can serve as a reliable quantitative tool for single-cell proteomic analysis.

## EXPERIMENTAL SECTION

### Cell Culture and Cell Counting

All cells were maintained in RPMI-1640 (Gibco), supplemented with 10% fetal bovine serum (FBS) and 1% penicillin streptococcus, and cultured in standard tissue culture conditions (37°C, 5% CO_2_). MDA-MB-231 was originally obtained from ATCC. Cell lines are routinely authenticated and tested for mycoplasma.

Parental MDA-MB-231 samples were prepared by seeding 1 × 10^6^ cells and incubating for 24 hours. Adherent cells were washed with PBS prior to being lifted with Cell Dissociation Buffer (Thermo Fisher Scientific). Lifted cells were re-suspended in 20 mL of culture medium and applied to a primed 15µM pluriStrainer^®^ (The Cell Separation Company) and the flow-through retained. 1 × 10^6^ cells were collected and washed twice in PBS. Following the final wash, supernatant was removed and the pellet was snap-frozen and stored at -80°C.

Drug-induced PACC samples were prepared by seeding 1 × 10^6^ MDA-MB-231 cells. After 24 hours incubation, cultures were treated with IC_50_ cisplatin or docetaxel for 72 hours. After 72 hours, surviving adherent cells were washed with PBS and lifted with Cell Dissociation Buffer (Thermo Fisher Scientific). Lifted cells were re-suspended in 20 mL of culture medium and filtered through primed 15µM pluriStrainer^®^ (The Cell Separation Company). The flow-through from each filter was discarded and the cells caught by the filter were harvested by flipping the filter upside-down and washing with 15 mL media. The PACC sample was pelleted at 1000 × g for 5 minutes, counted, and washed twice in PBS. Following the final wash, supernatant was removed and the pellet was snap-frozen and stored at -80°C.

### Sample Preparation

One million PACCs and parental MDA-MB-231 cancer cells (three samples from each cell type) were lysed in 60 µL lysis buffer containing 8M urea as described in CPTAC protocol^23^. Briefly, cell lysates were centrifugated at 16,000 × g for 12 min at 4°C and protein concentrations were determined by Pierce™BCA protein assay (Thermo). The samples were reduced by 6 mM dithiothreitol for 1h at 37 °C and then alkylated by 12 mM iodoacetamide for 45 min at room temperature in dark place. The samples were diluted to 2M urea concentration with 50 mM Tris buffer (pH 8.0). In the 2M urea buffer, the samples were digested with Lys-C (Wako) at 1 mAU: 10 mg enzyme to substrate ratio for 2h at room temperature, followed by the addition of trypsin (Promega) at the same ratio for overnight digestion at room temperature. After the digestion, the mixtures were acidified by 50% formic acid to get 1% formic acid as final concentration with pH<3. The digested peptides were desalted on C18 stage tips (3M) and dried with Speed-Vac (Thermo). Based on cell count and peptide yields (**S-Table 1**), each PACC sample was 0.2 ng peptides per cell and each parental MDA-MB-231 sample was about 0.05 ng peptides per cell. Starting from 1 µg aliquoted peptides, a serial of dilution was performed to obtain 100 cells, 10 cells and 1 cell populations for two different sizes of cells corresponding to 20ng, 2ng, and 0.2ng digested peptides for PACC, and 5ng, 0.5ng, and 0.05 ng digested peptides for parental cells. The dried peptides were redissolved in 3% acetonitrile with 0.1 % formic acid and used nanodrop to determine the peptide concentration. All the injections were spiked in 0.5 × iRT peptides (Biognosys) to calibrate the internal retention time.

### NanoLC-MS/MS Analysis

The aliquoted peptides equivalent to 100 cells, 10 cells, and a single cell of the PACCs and parental MDA-MB-231 cells were analyzed using two different LC gradient time, 15 minutes (min) and 120 min. All the samples, from 1 µg to 0.05 ng of peptides, were separated by EASY-nLC™ 1200 instrument (Thermo) with hand-packed analytical column (75 µm i.d. × 26.5 cm length packed with ReproSil-Pur 120 C18-AQ 1.9 µm beads) and Picofrit 10µm opening (New Objective). The column was heated to 50°C with Nanospray Flex™Ion Sources (Thermo). The elution flow rate was 200 nL/min with 0.1 % formic acid in 97 % H_2_O and 3 % CH_3_CN as buffer A, and 0.1 % formic acid in 90 % CH_3_CN and 10 % H_2_O as buffer B. Peptides were separated using 4-30% buffer B in 15 min gradient and 7-30% buffer B in 120 min gradient. All the samples were analyzed via Orbitrap Fusion Lumos Tribrid mass spectrometer (Thermo Fisher Scientific) and the parameters for the DIA method are as follows: resolution at 120,000, mass range of 350-1650 m/z, and maximum injection time of 60ms for MS1 scan; resolution at 30,000, HCD collision energy of 34%, mass range of 300-1600 m/z, and maximum injection time of 80ms for MS2 scan. For both MS1 and MS2, RF Lens 30% and normalized AGC Target 250% were applied. Total of 30 DIA raw files were acquired using two LC gradient settings from the aliquoted peptide samples of 100 cells, 10 cells and a single cell population of the PACC and the parental cancer cells (**S-Table 2**).

### DIA Data Analysis

The single-cell level DIA runs of global proteome were analyzed via library-free directDIA approach embedded in Spectronaut (version 14.10, Biognosys) with precursor and protein Qvalue cutoff at 1%. For the bulk sample analyses, five raw files (one from each of PACC samples and one from each of parental cell line samples) acquired from 1 µg injections were analyzed together in one directDIA search. For the single-cell analyses, the directDIA searches were conducted on five co-searching groups that each with different combination of raw files. Thus, each co-searching group generated an internal library containing different number of precursors (i.e., different library size). The five co-searching groups of single-cell analyses are as follows (**S-Table 3**): one single-cell raw file (271 precursor from 0.05ng peptides of a single MDA-MB-231 cell, referred to **GS-1r_M**; 2258 precursors from 0.2ng peptides of a single PACC cell, referred to as **GS-1r_P**), 10 raw files of all the single-cell injections from two LC gradients (2687 precursors, referred to as **GS-10r**), 16 raw files from the combination of **GS-10r** and six injections (two LC gradients) from 0.5ng peptides (10 cells) of MDA-MB-231 cells (5787 precursors, referred to as **GS-16r**), 20 raw files from the combination of **GS-10r** and ten-cell injections (two LC gradients) of PACC samples (16496 precursors, referred to as **GS-20r**), and all 30 raw files (47374 precursor, referred to as **G-30r**).

For the ten-cell analyses, the directDIA searches were conducted on five co-searching groups distinct from the co-searching groups of the single-cells as follows (**S-Table 4**): one ten-cell raw file (3803 precursor from 0.5ng peptides of a MDA-MB-231 cell sample, referred to as **G10c-1r_M**; 10756 precursors from 2ng peptides of a PACC sample, referred to as **G10c-1r_P**), 10 raw files of all the ten-cell injections from two LC gradients (17118 precursors, referred to as **G10c-10r**), 16 raw files by combining **G10c-10r** and all 100-cell injections of MDA-MB-231 cells (26191 precursors, referred to as **G10c-16r**), all 30 raw files (47374 precursor, referred to as **G-30r**), and 25 raw files as a combination of **G10c-10r**, all 100-cell injections of PACC samples and five 1µg peptide injections (88012 precursors, referred to as **G10c-25r**).

The PTM analyses were conducted by searching global DIA data (nano-gram and single-cell levels) against the pre-built PTM spectral libraries.

### PTM Spectral Library Generation

To analyze PTMs, the spectral libraries for **Fig. 4a** were generated from patient-derived xenograft (PDX) samples, phosphopeptide spectral library was built using IMAC-enriched phosphorylation data of PDX samples (followed CPTAC standard protocol^24^), acetylation and ubiquitination spectral libraries were constructed using the data of antibody-enriched PDX samples^25,26^. All PTM spectral libraries from PDX samples were built by single DIA run, individually. Additionally, three phosphopeptide spectral libraries with different sizes (∼42,000 precursors, ∼87,000 precursors and ∼142,000 precursors) were built using IMAC-enriched phosphorylation data of tumor tissues from clear cell renal cell carcinoma (ccRCC) tissues^24^. All the PTM libraries were built by using Pulser embedded in Spectronaut.

## RESULTS AND DISSCUSSION

### LC Gradient Time for Protein Identification at Single-Cell Levels

In general, an optimal protein identification can be achieved using 1 µg peptides for DIA-MS analysis in combination with 120-min LC gradient for large cell population (⩾20,000 cells) and bulk cell samples^27^. However, such approach may not be ideal for small cell population. Therefore, we conducted a comparative analysis on different LC gradient settings for peptides equivalent to hundred-cell, ten-cell, and single-cell levels. We first compared two LC gradient settings for a single cell (0.05 ng peptides), 10 cells (0.5 ng) and 100 cells (5 ng) from MDA-MB-231 cell samples by computing the identification ratio of 15-min LC gradient to 120-min LC gradient based on the average number of identified proteins. We found the number of protein identification rate was more in 15-min LC gradient for single-cell and 10-cell levels. As shown in **Fig. 1a**, the protein identification ratios of 15-min to 120-min were more than 1 for single and ten MDA-MB-231 cells. We observed similar result for the single cisplatin treated PACC cell (0.2ng of peptides) (**Fig. 1b**), where the ratio of 15-min to 120-min was also more than 1, indicating that less proteins were identified from 120-min LC gradient and an improvement in overall protein identification using a short LC gradient time at single-cell level of peptide injection amount < 2ng. Therefore, we chose 15 minutes as our optimal LC gradient setting for single-cell level DIA analysis.

**Fig. 1.**
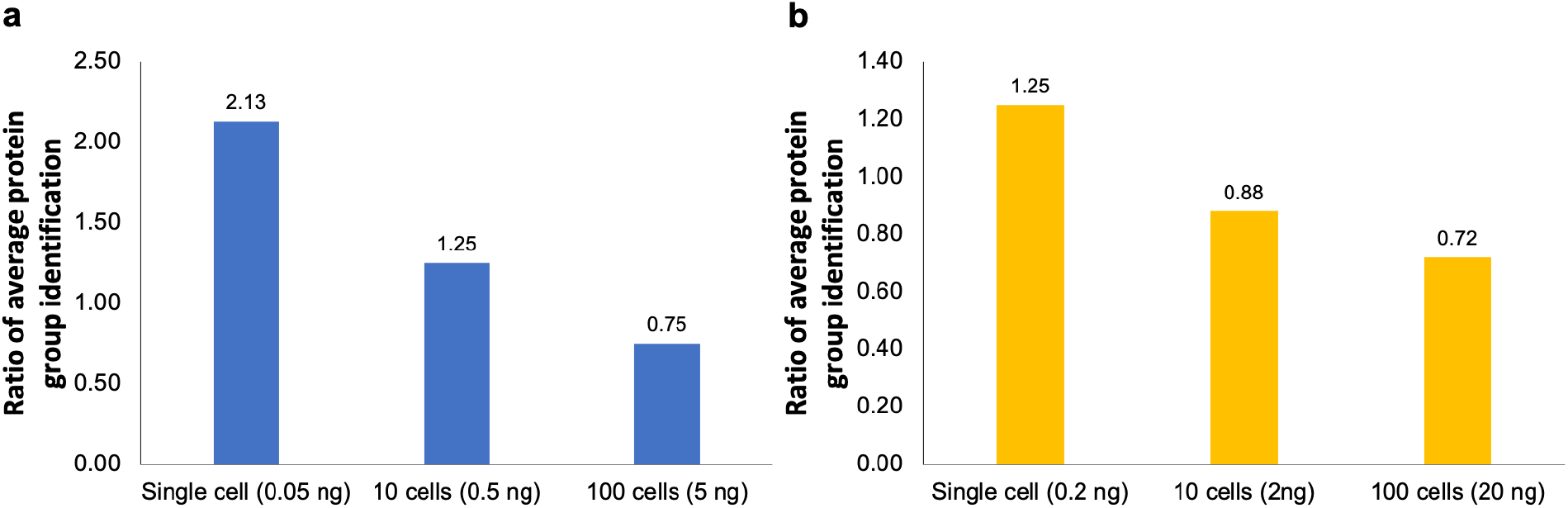
Comparison of 15-min vs 120-min liquid chromatography (LC) gradient for proteomic analysis of limited number of MDA and PACC cells. Protein identification ratios of 15-min LC gradient to 120-min LC gradient for DIA analyses on (a) MDA-MB-231 cells and (b) cisplatin treated PACCs.

### Evaluation of Global Proteomic Analysis at Single-Cell Level

Besides investigating the suitable LC gradient for acquiring DIA-MS data at single-cell level, the search space of single-cell DIA data (i.e., the size of internal library generated during directDIA search) was also evaluated among the established co-searching groups. The DIA data of MDA-MB-231 cell samples (0.05 ng peptides equivalent to the single cell level (0.05 ng), searching GS-1r_M (total of 271 peptide precursors from the raw file of a single MDA-MB-231 cell), and 126 protein groups were identified (**Fig. 2a**). As the size of the internal library increased to 5787 precursors (i.e., GS-16r), we observed the highest peptide and protein coverage for the single MDA-MB-231 cancer cell. Total of 1093 peptide precursors and 406 protein groups were identified using GS-16r (**Fig. 2a**), corresponding to gains of 303% and 222% at peptide and protein levels compared to the results obtained by using GS-1r_M only. For cisplatin-treated PACC at single-cell level (0.2 ng of peptides), 621 protein groups were identified using the directDIA approach to search against the internal library of GS-1r_P (2258 peptide precursors) (**Fig. 2b**). Moreover, we found that using co-searching group of GS-20r (16496 precursors) yielded the best identification, where 6153 peptide precursors and 1530 protein groups were identified (**Fig. 2b**). By using GS-20r, 172% and 146 % gains at peptide precursor and protein levels relative to using GS-1r_P, respectively. Of note, the number of identified proteins/peptides was not necessarily increased as the search space expanded. As shown in **Fig. 2c**, the best precursor identification is within the range of 2-to 4-fold difference between the total number of precursors in an internal library and total number of precursors detected in the sample of interest. Our results suggested that the internal library size was critical to protein identification at single-cell and sub-nano-gram levels via DIA-MS approach.

**Fig. 2.**
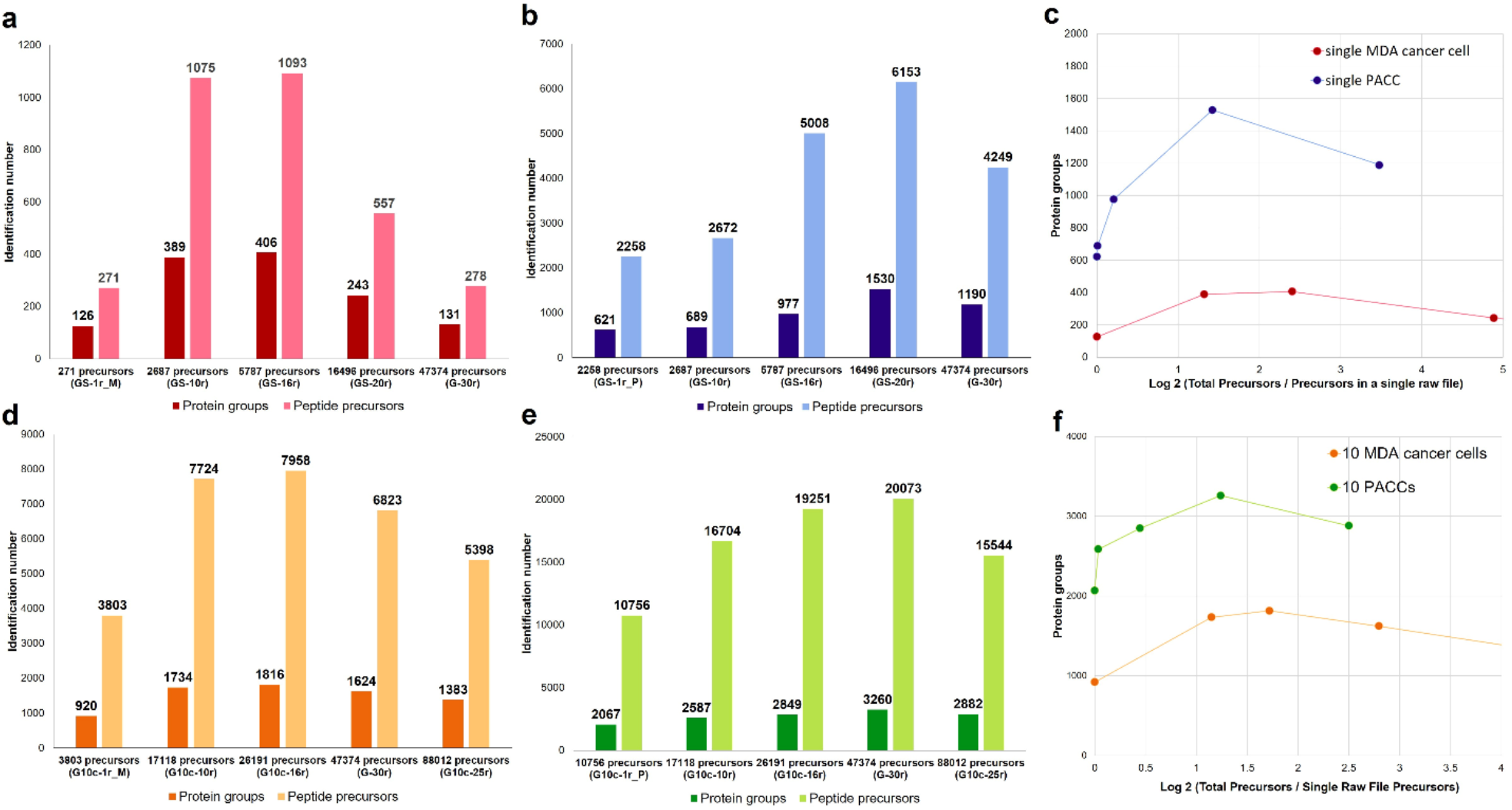
Evaluation of different co-searching groups (internal libraries with different numbers of precursors) generated during directDIA search for single-cell and ten-cell proteomic DIA analysis. (a) Numbers of peptides and proteins identified from different co-searching groups at the single MDA-MB-231 cell level. (b) Numbers of peptides and proteins identified from different co-searching groups at the single PACC level. (c) The single-cell level protein identification towards the ratio of total precursors comparing to single cell level precursor identifications with matched co-searching precursors. (d) Numbers of peptides and proteins identified from different co-searching groups at the ten MDA-MB-231 cell level. (e) Numbers of peptides and proteins identified from different co-searching groups at the ten PACC level. (f) The ten-cell level protein identification towards the ratio of total precursors comparing to ten-cell level precursor identifications with matched co-searching precursors.

Furthermore, we evaluated the co-searching methods at ten-cell level. We identified 1816 protein groups (**Fig. 2d**) and 3260 protein groups (**Fig. 2e**) from 10 MDA-MB-231 cells and 10 PACCs, respectively. Similarly, the best peptide precursor and protein identifications were also fallen into the 2-to 4-fold changes between the internal library size and detected precursors (**Fig. 2f**). In addition, we observed the optimal protein identification via directDIA search for the single-cell and ten-cell injections when co-searched with injected peptide amount that were about10-fold difference within the similar samples (**S-Table 5**). Overall, it was essential to use a co-searching internal library generated from similar samples during the directDIA search to enhance protein identification at the single-cell and nano-gram levels.

### Reproducibility on Single-Cell Level Proteomic Analysis Using DIA

After evaluation of LC gradient and size of internal library for DIA analysis at single-cell level, we investigated the inter-person reproducibility using our established workflow. Two sets of samples, each contained cisplatin- and docetaxel-treated PACC and parental MDA-MB-231 cancer cells, were prepared and analyzed a month apart by two researchers following the procedures stated in Materials and Methods section. Here, we use the cisplatin treated PACC sample and one MDA-MB-231 sample to demonstrate the reproducibility of our DIA method. We observed pairwise Spearman correlation >0.80 for the MDA-MB-231 cell sample between the two sets at single-cell, ten-cell, and hundred-cell levels (**Fig 3a**). We found similar results for cisplatin treated PACC samples, where Spearman correlation ⩾0.83 were observed in three different cell populations (**Fig 3b**). Taken together, these results demonstrated that our optimized DIA workflow provided robust quantitative global proteome profiling for single-cell DIA analysis, and larger cell population further improved the reproducibility.

**Fig. 3.**
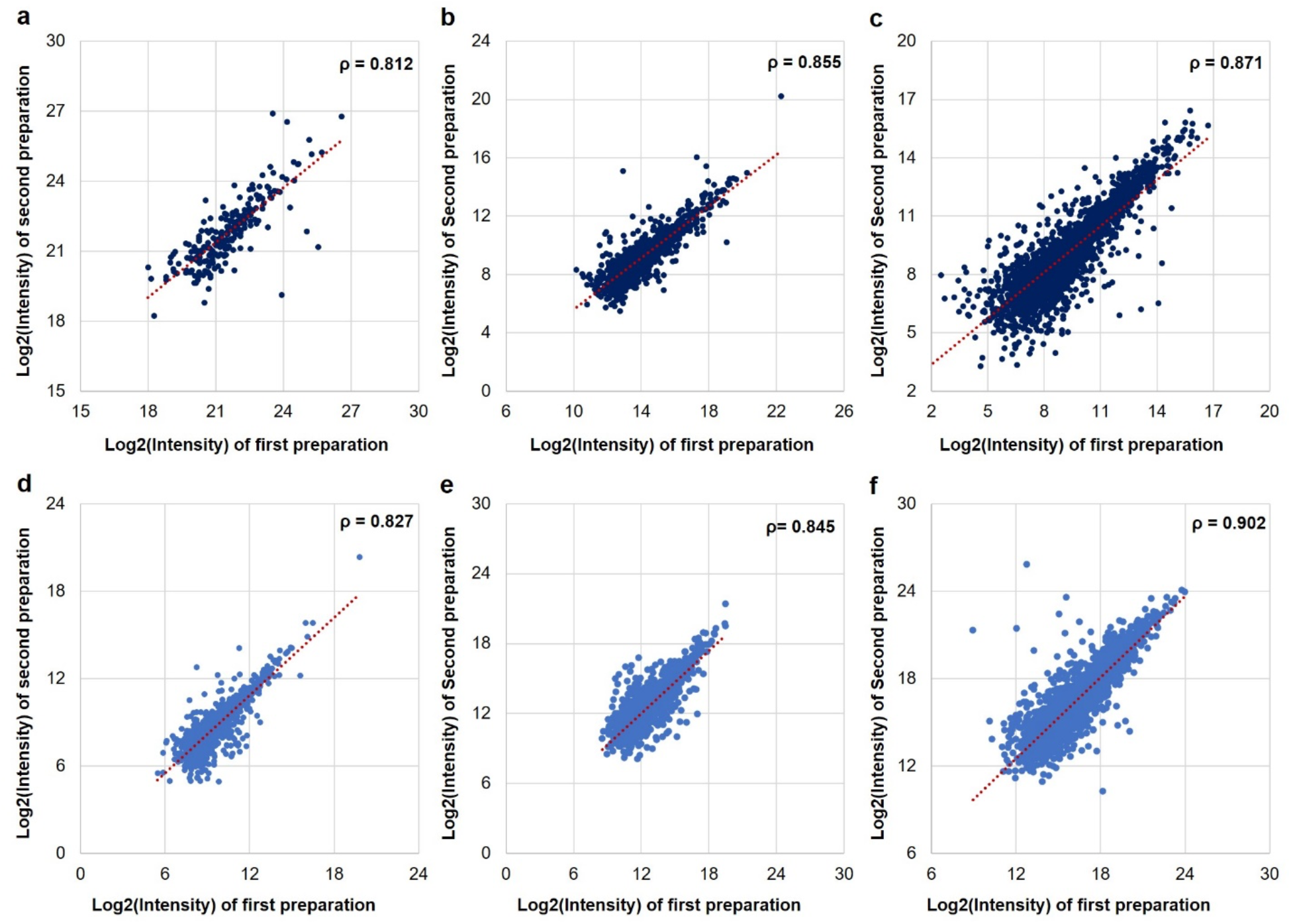
Reproducibility of the DIA-MS runs where the sample preparations were done month apart by two different researchers. Pairwise Spearman correlation of (a) protein intensities between two preparations and DIA-MS analyses of MDA-MB-231 cells and (b) protein intensities between two preparations and DIA-MS analyses of cisplatin treated PACCs.

### PTM Analyses for Nano-Gram Levels of Peptides without Enrichment

Protein modifications are important for the regulation of various protein activities and cellular signaling events, and alternation in PTMs are associated with many diseases, including cancer^28^. When conducting MS-based PTM analysis, PTM enrichment is an essential procedure; however, there is very limited report of nano-gram/single-cell level of enrichment strategies for MS analysis. Thus, we established an alternative approach for PTM analysis at such level by utilizing global proteomic DIA data and spectral libraries built from bulk samples. Unlike DDA-MS, DIA-MS allows that all the peptide precursors are co-fragmented within a selected m/z range to produce comprehensive MS2 spectra. The information of modified peptides should be retained in the global data even without PTM enrichment. Therefore, we were able to directly identify PTMs from the nano-gram level (i.e., 100 cells) of global proteomic DIA data using customized PTM spectral libraries for phosphorylation, acetylation, and ubiquitination.

We firstly explored the possibility of identifying phosphorylation, acetylation, and ubiquitination from global DIA data of PDX samples at nano-gram level (**Fig. 4a**). We identified 72 phosphorylated peptides, 35 acetylated peptides, and 99 ubiquitinated peptides from 100 PACCs, indicating the possibility of finding PTMs without using enrichment.

**Fig. 4.**
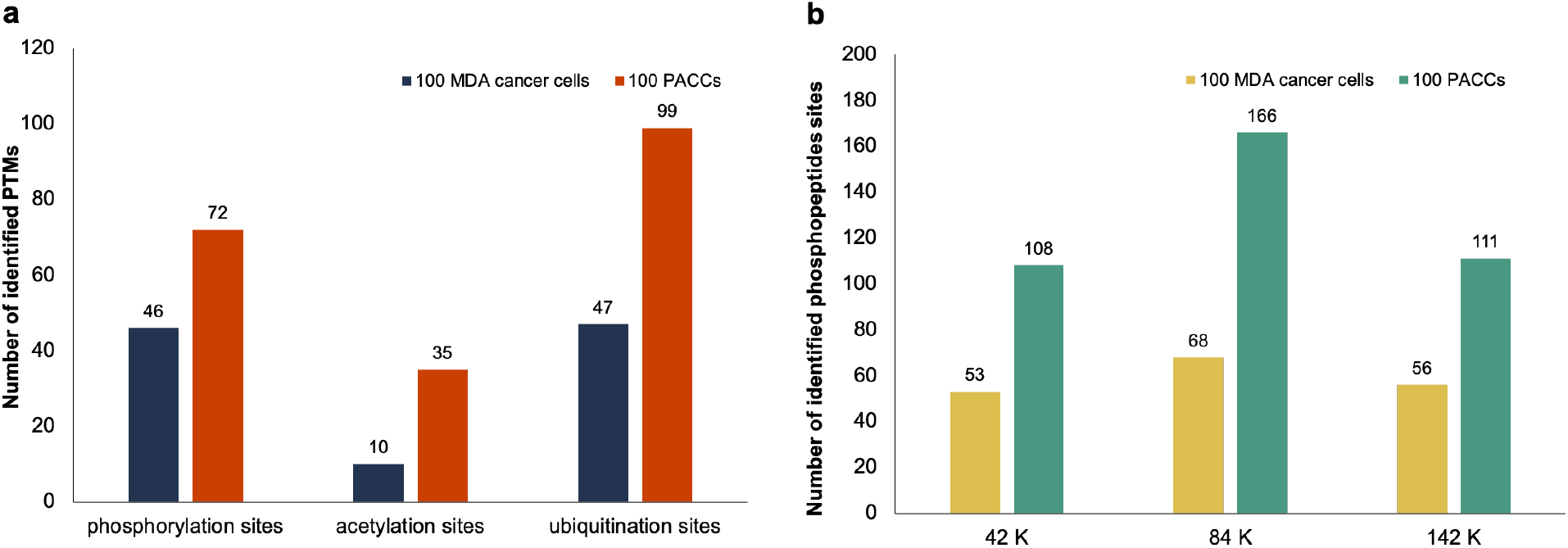
Identifications of PTM peptides/sites from global DIA-MS runs. (a) The numbers of identified PTM peptides/sites from 100 cells via the spectral libraries constructed from PTM-enriched DIA data of PDX samples. (b) Evaluation of phosphopeptide spectral library size towards phosphorylation site identification from the spectral libraries built using phopshopeptide-enriched from ccRCC samples.

We further evaluated the association between PTM spectral library size and PTM identification by examining the alteration in phosphopeptide identification rate from the nano-gram level of global DIA data, since a large collection of phosphopeptide-enriched DDA and DIA raw files from CPTAC study^22^ allowed the construction of spectral libraries with various sizes ranging from ∼42K to ∼141K precursors. As shown in **Fig. 4b**, among the three phosphopeptide spectral libraries, the library containing ∼84K precursors contributes to the highest identification number for 100 MDA-MB-231 cells (5 ng of peptides) and 100 PACCs (20 ng of peptides) of which 68 and 166 phopshopeptides with localized sites are identified, respectively. These results suggested that PTM analysis of nano-gram scale could be achieved by utilizing global DIA data along with a suitable PTM library built from bulk samples.

### Application of Single-Cell Level DIA Approach to the Drug Resistant Cancer Cell Study

To investigate whether the difference in cell size affected identification and protein expression patterns, we conducted a comparative analysis between PACCs (large cells) and MDA-MB-231 cells (smaller cells) at 1 µg peptide injection and single-cell level of peptide injection. We observed 98.5% of overlap in protein identification between cisplatin treated PACC and MDA-MB-231 samples (**Fig. 5a**), suggesting that they shared similar proteome profile, regardless of cell size. At single-cell level, we examined the identified protein groups from the co-searching via all single-cell raw files (i.e., GS-10r, **Fig. 2a**). We observed 388 protein groups identified in the single MDA-MB-231 cell and 688 proteins were identified from the cisplatin-treated PACC at single cell level (**Fig. 5b**).

**Fig. 5.**
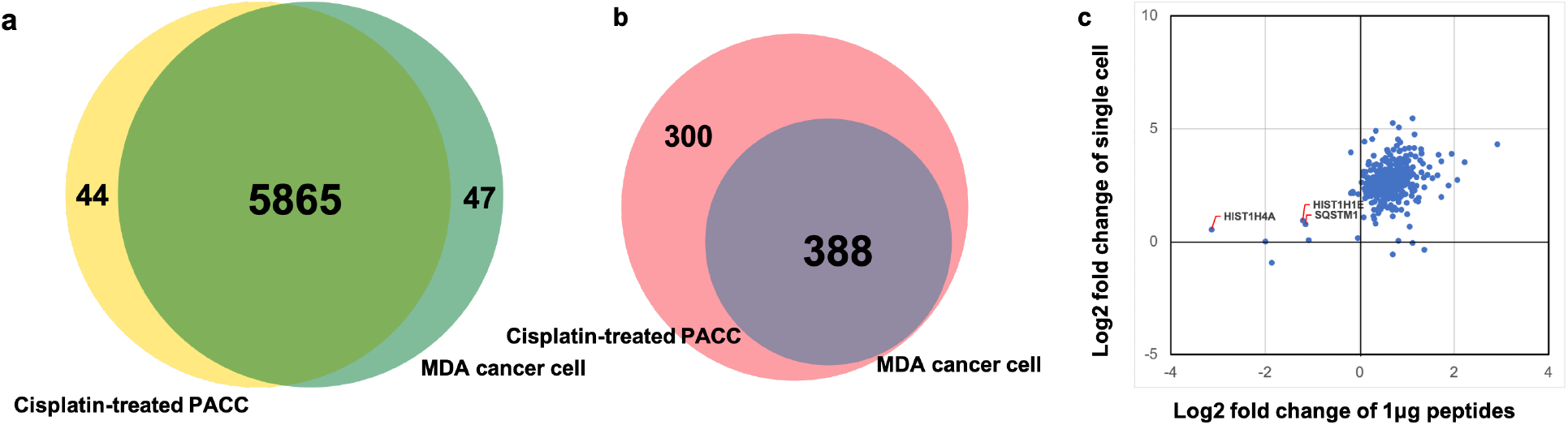
Global and single-cell level comparison of quantitative proteomics from cisplatin resistant PACC and MDA-MB-231 cells. (a) Protein identification comparison using 1 µg peptide amount injections. (b) Protein identification comparison at single-cell level. (c) Quantitative analysis between single-cell level and 1 µg of peptides in terms of the comparison in Log2 fold changes between MDA-MB-231 cells and PACC cells.

Although more proteins were identified using 1 µg peptide injections, we speculated that single-cell proteomic analysis could capture the real protein expression changes comparing to bulk proteomic analysis, which would assist in the study of cellular heterogeneity between PACC and parental MDA-MB-231 cells. We compared the protein fold changes as shown in **Fig. 5c**. At single-cell level, we found majority of the proteins showing higher expression in cisplatin resistant PACC relative to the parental MDA-MB-231 cell with Log2 fold changes ranged between 2 to 5, indicating protein copy number increase for these proteins after treating MDA-MB-231 cells with cisplatin and transitioning to a PACC state, except ubiquitin-binding autophagy associated protein (e.g., SQSTM1) and histone proteins (e.g., HIST1H4A and HIST1H1E) displayed similar expression profiles between PACCs and control cancer cells. In contrast, in bulk analysis of the MDA-MB-231 and cisplatin-treated PACCs, most of the proteins showed similar abundances at the same injection amount (1.0 µg) level; however, we found >2-fold decrease in PACC compared to MDA-MB-231 cancer cells for SQSTM1, HIST1H4A and HIST1H1E. If we only run the samples at 1µg peptide level, we could only observe the decrease in SQSTM1, HIST1H4A and HIST1H1E protein intensities in PACCs. However, at the single-cell level analyses, we noticed these proteins maintaining in the same expression levels in both PACCs and parental MDA-MB-231 cells. The same expression level of SQSTM1 suggested PACCs and MDA-MB-231 cells had the similar metabolic activity^29^, which may be not enough for a PACC to undergo depolyploidization to transition to typical MDA-MB-231 cells. As lack of multi-copy of histone proteins leading to cell cycle elongation^30,31^ and PACCs are also unable to divide to normal MDA-MB-231 cancer cells. Taken together, quantitative analysis of single-cell proteome via DIA approach can benefit our understanding of cellular heterogeneity and provide more accurate protein expression profiles which may be misinterpreted at bulk population.

## CONCLUSIONS

Single-cell proteomic analysis provides insights into cellular heterogeneity allowing the characterization of cellular microenvironment, whereas proteomic analysis using bulk samples only captures a population average hindering our understanding of the diversity in cellular functions. In this study, we established and optimized a single-cell proteomic analysis workflow which utilized DIA-MS and directDIA method to analyze the global proteome of the MDA-MB-231 cancer cells and the matched drug resistant PACC cells. We first systematically evaluated aliquoted peptide samples from a single cell to 100 cell levels (0.05 ng to 20 ng) of the two cell types under different LC gradient settings. We found that 120-min LC gradient was more suitable for peptide injection amount > 2 ng (∼10 PACCs), whereas 15-min LC gradient produced was more appropriate for the injection amount < 2 ng (∼10 PACCs). By applying and investigating directDIA search method using different co-searching groups (i.e., internal libraries), we observed approximately a 4-fold difference between the internal library size and total number of detected precursors of a DIA raw file produced the highest protein identification rate with good reproducibility. Of note, 1500-3000 proteins were identified from 10-140 mammalian cells (equaled to 0.5-7 ng of peptides) by using narrow-bore columns (i.d. 30µm or 20 µm) coupled with low flow rate separation^32^, where <2000 proteins were identified from 2 ng aliquoted peptide samples by using a special LC system^33,34^. Nevertheless, using our optimized workflow, 2ng (∼10 PACCs) of peptides allowed an identification of ≥3200 proteins and 1500 proteins were identified from a single PACC cell (∼0.2 ng of peptides) even with normal bore column (i.d. 75µm) and normal LC system. Furthermore, identification of PTMs at nano-gram/single-cell level without any PTM enrichment was achieved by directly searching the nano-gram level of global DIA data against pre-generated PTM libraries. Additionally, we were able to detect the cellular heterogeneity between PACCs and their parental MDA-MB-231 cells at single-cell level using our established workflow. In summary, we developed a novel approach to study small cell population, including single cell, by using DIA-MS coupled with a short LC gradient and the directDIA search with an internal library with an appropriate size, which can support quantitative single-cell proteomic and PTM analyses at high-throughput.

## ACKNOWLEDGMENTS

This work was supported by fundings from: National Cancer Institute, the Clinical Proteomic Tumor Analysis Consortium (CPTAC, U24CA210985); National Cancer Institute, Early Detection Research Network (EDRN, U01CA152813); US Department of Defense CDMRP/PCRP (W81XWH-20-10353), the Patrick C. Walsh Prostate Cancer Research Fund and the Prostate Cancer Foundation to SR Amend; and NCI grants U54CA143803, CA163124, CA093900, and CA143055, and the Prostate Cancer Foundation to KJ Pienta.

## AUTHOR CONTRIBUTIONS

H.Z., Y.W., T.M.L., L-W.C., M.D.K., S.R.A., and K.J.P. conceived and designed the experiments; H.Z., Y.W. and T.M.L. interpreted the results; M.D.K. performed the cell culture experiments; Y.W. and L-J.C. performed the experiments of mass spectrometry part; Y.X. prepared the PTM libraries; Y.W., T.M.L. and H.Z. analyzed data and prepared figures; Y.W. drafted the manuscript; T.M.L., H.Z., M.D.K., K.J.P, and S.R.A. edited the manuscript; H.Z. oversaw the execution of the project.

## REFERENCES

1. Macaulay, I. C.; Ponting, C. P.; Voet, T.; Single-Cell Multiomics: Multiple Measurements from Single Cells. Trends Genet., 32, 155–168 (2017).

2. Lee, J.; Hyeon, D. Y.; Hwang, D.; Single-Cell multiomics: technologies and data analysis methods. Exp. Mol. Med., 52, 1428–1442 (2020).

3. De Rosa, S. C.; Herzenberg, L. A.; Herzenberg, L. A.; Roederer, M.; 11-color, 13-parameter flow cytometry: Identification of human naive T cells by phenotype, function, and T-cell receptor diversity. Nat. Medicine, 7, 245–248 (2001).

4. Perez, O. D.; Nolan, G. P.; Simultaneous measurement of multiple active kinase states using polychromatic flow cytometry. Nat. Biotechnol., 20, 155–162 (2002).

5. Bandura, D. R. et al. Mass cytometry: technique for real time single cell multitarget immunoassaybased on inductively coupled plasma time-of-flight mass spectrometry. Anal. Chem., 81, 6813–6822, (2009).

6. Hughes, A. J.; Spelke, D. P.; Xu, Z.; Kang, C.; Schaffer, D. V.; Herr, A. E.; Single-cell western blotting. Nat. Methods, 11, 749–755 (2014).

7. Budnik, B.; Levy, E.; Harmange, G.; Slavov, N.; SCoPE-MS: mass spectrometry of single mammalian cells quantifies proteome heterogeneity during cell differentiation. Genome Biol., 19, 161, (2018).

8. Ctortecka, C.; Mechtler, K.; The rise of single-cell proteomics. Anal. Sci. Adv., doi.org/10.1002/ansa.202000152 (2021)

9. Cheung, T. K.; Lee, C.; Bayer, F. P.; McCoy, A.; Kuster, B.; Rose, C. M.; Defining the carrier proteome limit for single-cell proteomics. Nat. Methods, 18, 76–83 (2021).

10. Gillet, L. C.; Navarro, P.; Tate, S.; Röst, H.; Selevsek, N.; Reiter, L.; Bonner, R.; Aebersold, R.; Targeted Data Extraction of the MS/MS Spectra Generated by Data-independent Acquisition: A New Concept for Consistent and Accurate Proteome Analysis. Mol. Cell Proteomics, 11:O111.016717 (2012)

11. Collins, B. C. et al. Multi-laboratory assessment of reproducibility, qualitative and quantitative performance of SWATH-mass spectrometry. Nat. Commun., 8, 291 (2017).

12. Muntel, J.; Kirkpatrick, J.; Bruderer, R.; Huang, T.; Vitek, O.; Ori, A.; Reiter, L.; Comparison of Protein Quantification in a Complex Background by DIA and TMT Workflows with Fixed Instrument Time. J. Proteome Res., 18, 1340–1351 (2019).

13. Thomas, S. N.; Friedrich, B.; Schanubelt, M.; Chan, D. W.; Zhang, H.; Aebersold, R.; Orthogonal Proteomic Platforms and Their Implications for the Stable Classification of High-Grade Serous Ovarian Cancer Subtypes. iScience, 23, 101079 (2020).

14. Navarro, P. et al. A multicenter study benchmarks software tools for label-free proteome quantification. Nat. Biotech., 34, 1130–1136 (2016).

15. Zhang, F.; Ge, W.; Ruan, G.; Cai, X.; Guo, T.; Data-Independent Acquisition Mass Spectrometry-Based Proteomics and Software Tools: A Glimpse in 2020. Proteomics, 20, 1900276 (2020).

16. Bruderer R. et al. Extending the limits of quantitative proteome profiling with data-independent acquisition and application to acetaminophen-treated three-dimensional liver microtissues. Mol Cell Proteomics, 14, 1400–10 (2015).

17. Tsou, C.; Avtonomov, D.; Larsen, B.; Tucholska, M.; Choi, H.; Gingras, A.; Nesvizhskii, A. I.; DIA-Umpire: comprehensive computational framework for data-independent acquisition proteomics. Nat. Methods, 12, 258–64 (2015).

18. Parker, S. J.; Venkatraman, V.; Van Eyk, J. E.; Effect of peptide assay library size and composition in targeted data-independent acquisition-MS analyses. Proteomics, 16, 2221–2237 (2016)

19. Wu, J. X.; Song, X.; Pascovici, D.; Zaw, T.; Care, N.; Krisp, C.; Molloy, M. P.; SWATH Mass Spectrometry Performance Using Extended Peptide MS/MS Assay Libraries. Mol. Cell Proteomics, 15, 2501–2514 (2016).

20. Barkovits, K. et al. Reproducibility, Specificity and Accuracy of Relative Quantification Using Spectral Library-based Data-independent Acquisition. Mol. Cell Proteomics, 19, 181–197 (2020).

21. Muntel, J.; Gandhi, T.; Verbeke, L.; Bernhardt, O. M.; Treiber, T.; Bruderer, R.; Reiter, L.; Surpassing 10 000 identified and quantified proteins in a single run by optimizing current LC-MS instrumentation and data analysis strategy. Mol. Omics, 15, 348–360 (2019).

22. Pienta, K. J.; Hammarlund, E. U.; Axelrod, R.; Brown, J. S.; Amend, S. R.; Poly-aneuploid cancer cells promote evolvability, generating lethal cancer. Evolut. Appl., 13, 1626–1634 (2020).

23. Mertins, P. et al. Reproducible workflow for multiplexed deep-scale proteome and phosphoproteome analysis of tumor tissues by liquid chromatography–mass spectrometry. Nat. Protocals, 13, 1632–1661 (2018).

24. Clark, D. J. et al. Integrated Proteogenomic Characterization of Clear Cell Renal Cell Carcinoma. Cell, 179, 964–983 (2019).

25. Wang, L. et al. Proteogenomic and metabolomic characterization of human glioblastoma. Cancer Cell, 39, 509-528.e20 (2020).

26. Udeshi, N. D. et al. Rapid and deep-scale ubiquitylation profiling for biology and translational research. Nat. Commun., 11, 539 (2020).

27. Cho, K. et al. Deep Proteomics Using Two Dimensional Data Independent Acquisition Mass Spectrometry. Anal. Chem., 96, 4217–4225 (2020).

28. Chen, L.; Liu, S.; Tao, Y.; Regulating tumor suppressor genes: post-translational modifications. Signal Transduct. Target. Ther., 5, 90 (2020).

29. Sánchez-Martín, P.; Komatsu, M.; p62/SQSTM1 – steering the cell through health and disease. J. Cell Sci., 131, jcs222836 (2018).

30. Marzluff, W. F.; Wagner, E. J.; Duronio, R. J.; Metabolism and regulation of canonical histone mRNAs: life without a poly(A) tail. Nat Rev Genet., 9, 843–854 (2008).

31. Günesdogan, U.; Jäckle, H.; Herzig, A.; Histone supply regulates S phase timing and cell cycle progression. eLife, 3, e02443 (2014).

32. Zhu, Y. et al. Nanodroplet processing platform for deep and quantitative proteome profiling of 10–100 mammalian cells. Nat. Commun., 9, 882 (2018).

33. Brunner, A. et al. Ultra-high sensitivity mass spectrometry quantifies single-cell proteome changes upon perturbation, bioRxiv, doi: 10.1101/2020.12.22.423933 (2021)

34. Cong, Y. et al. Improved Single-Cell Proteome Coverage Using Narrow-Bore Packed NanoLC Columns and Ultrasensitive Mass Spectrometry. Anal. Chem., 92, 2665–2671 (2020)

